# Ancestral Hydrocarbon Metabolism Enables PET Degradation by a Natural Bacterial Consortium

**DOI:** 10.64898/2026.03.18.709718

**Authors:** Sabrina Edwards, Danny Rice, Patricio Palomino, Irene Newton, Jay L. Mellies

## Abstract

Plastic biodegradation in natural environments is increasingly recognized as a multi-organism process, yet the mechanisms enabling coordinated depolymerization and metabolism of polyethylene terephthalate (PET) remain poorly understood. Previously, we demonstrated that a full consortium containing three *Pseudomonas* and two *Bacillus* strains isolated from hydrocarbon-rich coastal soils of Galveston Bay, Texas, can synergistically depolymerize PET plastic and utilize it as a sole carbon source, a capacity not observed in individual isolates. In this report, using integrated comparative genomics, proteomics, and chemical analyses, we show that PET degradation in this system reflects exaptation of hydrocarbon metabolism reinforced by metabolic division of labor. Within this naturally occurring consortium, *Bacillus* strains persist under environmental stress, establish biofilms, and perform essential secondary hydrolysis, while *Pseudomonas* strains catabolize aromatic monomers and buffer oxidative stress. Genes supporting these functions are enriched within the accessory genomes of the consortium strains, indicating consortium-enriched horizontal gene transfer (HGT). In addition to the canonical two-step hydrolytic pathway well documented in PET biodegradation, we identify a secondary methylation-and redox-associated process, mechanisms where the full consortium acts on the oligomer mono(2-hydroxyethyl) terephthalate (MHET), yielding nearly complete conversion to terephthalic acid (TPA) and methylated MHET (MMHET). Together, these findings demonstrate how cooperation and competition within consortia facilitate targeted gene exchange, enabling emergent plastic biodegradation in natural microbial communities.

**IMPORTANCE:** Environmental plastic degradation is rarely accomplished by a single organism, yet the microbial mechanisms enabling community-level PET plastic breakdown remain poorly understood. This study shows that a bacterial consortium isolated from crude petroleum-contaminated beaches biodegrades PET through exaptation of ancestral hydrocarbon pathways, metabolic division of labor, and targeted gene exchange rather than specialized PET-specific metabolic pathways. *Pseudomonas* strains initiate PET cleavage, while stress-tolerant *Bacillus* strains persist long enough to clear inhibitory intermediates and enable downstream aromatic and diol metabolism. PET degradation is observed to be an emergent property of ecological interactions and distant evolutionary history. These findings provide a community-level model for understanding how natural microbial communities may adapt to novel anthropogenic substrates such as synthetic polymers, sustaining prolonged biodegradation.

## INTRODUCTION

Galveston Bay, Texas, hosts extensive natural oil seeps where crude petroleum has migrated into coastal soils for millennia and has been a hotspot of anthropogenic oil spills (1–3). As a result, these chronically hydrocarbon-exposed environments are dominated by hydrocarbonoclastic bacteria (HCB), including the genera *Pseudomonas*, *Acinetobacter*, *Rhodococcus*, and *Bacillus* (4–8). Over evolutionary timescales, these taxa have refined pathways for utilizing petroleum-derived compounds(6–13). Building on this ecological context, we previously isolated a microbial consortium, from Galveston Bay, composed of three *Pseudomonas* and two *Bacillus* strains that synergistically degrade both polyethylene terephthalate (PET) and polyethylene (PE) plastics(14,15) Because synthetic plastics are synthesized from petroleum-derived monomers, we hypothesized that hydrocarbon-adapted bacteria within this consortium may be evolutionarily primed to exploit plastic-associated substrates through pre-existing hydrocarbon metabolic pathways.

Polyethylene terephthalate (PET), the most widely used packaging and textile polymer, is assembled from terephthalic acid (TPA) and ethylene glycol (EG), while polyethylene (PE) consists of long hydrocarbon chains. Although these monomers are readily metabolized by HCB, such as this described consortium, once polymerized, these compounds form highly durable plastic polymer that persist in the environment, polluting aquatic and terrestrial ecosystems(16–20). When plastics enter the environment, abiotic weathering (UV, oxidation, mechanical stress) gradually fractures polymer chains, introducing hydrophilic groups, increasing surface reactivity, and enabling microbial colonization(21–25). These changes increase bioavailability and allow microbes to form complex biofilms and secrete extracellular enzymes that fragment polymers into soluble oligomers and monomers, which can be converted to metabolic intermediates and subsequently funneled back into pre-existing metabolic pathways(26,27).

To date, researchers have identified a large and growing catalog of individual polyethylene terephthalate (PET)–degrading enzymes. These include PETase from *Ideonella sakaiensis*, isolated from e-waste–associated environments, and highly active thermophilic cutinases derived from biopolymer-rich sites such as leaf compost (*e.g*., leaf compost cutinase, LCC) (28–30), which can either partially cleave PET to oligomers bis-(2-hydroxyethyl) terephthalate (BHET) and mono-(2-hydroxyethyl) terephthalate (MHET), or fully to the original monomers, TPA and EG. While these enzymes demonstrate that PET hydrolysis is biochemically feasible for biotechnology recycling efforts, environmental plastic degradation is rarely attributable to a single enzyme. Instead, increasing evidence suggests that PET breakdown in natural systems reflects coordinated activity within microbial consortia, driven by metabolic specialization across community members (30–33). Despite this knowledge, the mechanisms enabling cooperative and synergistic PET degradation and assimilation in environmental consortia remain poorly resolved. It remains unclear whether these biodegradation capabilities arise primarily through horizontal gene transfer, gradual sequence refinement, or exaptation of pre-existing hydrocarbon metabolic pathways (*i.e*. features that originally evolved for petroleum degradation but may be co-opted for plastic-associated substrates). Although PET biodegradation has been partially characterized in several systems, the evolutionary basis of PET degradation in natural microbial communities is still not well understood (29,34).

Previous pangenomic and transcriptomic analyses of the five-member consortium initially revealed that these isolates shared significant overlaps in accessory genomes compared to non-PET degraders, including genes implicated in the degradation of multiple bio- and synthetic-plastics, like numerous hydrolases and oxidoreductases (35). Given that this consortium may have been evolutionarily “primed” to exploit plastic-associated substrates due to their hydrocarbon-rich origins, we sought to resolve how these functions are distributed and coordinated during PET degradation using horizontal gene transfer (HGT) (36) analysis and functional characterization of secreted proteomes during growth on PET. These datasets revealed significant inter-genera genetic sharing, and complementary roles in extracellular depolymerization, redox handling, and downstream processing of PET-derived intermediates, a pattern that was further supported by the detection of methylated mono(2-hydroxyethyl) terephthalate (MMHET) during PET incubation.

Our results show that PET degradation is not the product of a single enzyme or pathway, but community-level adaptation. This *Bacillus-Pseudomonas* consortium exhibits synergistic PET degradation mediated through a cooperative division of labor, as complete PET mineralization occurs when all strains are present. When strains are absent, MHET accumulates, which can inhibit further depolymerization, indicating true metabolic interdependence. Notably, MHET catabolism does not exclusively follow canonical hydrolytic fates, suggesting the emergence of alternative, environmentally relevant chemical transformations during PET depolymerization. In this study, we examine evidence that adaptation to plastic as a substrate is consortia-based and distributed across complementary physiological roles, where survival and carbon assimilation depend on cooperation, stress buffering, and coordinated metabolic activity.

## MATERIALS AND METHODS

### Bacterial Strains and Bioinformatics

The five bacterial strains comprising the consortium used in this study were previously obtained from petroleum-polluted soils in Galveston Bay, Texas(37). Genomes were sequenced using hybrid Illumina/Nanopore technology and assembled with Unicycler v0.5.0 (BioProject <u>PRJNA517285</u>). Strains include *Bacillus cereus* 9.1 and *B. thuringiensis* 9.1 (SAMN41232461, - 58) and *Pseudomonas* spp. strains 9.2, 10, and 13.2 (SAMN41232459, -60, -62). IMG/MER annotations were used for feature detection, while NCBI GenBank IDs served as the primary identifiers. Multi-omic data (transcriptomics, proteomics, HGT) were standardized by mapping sequences to NCBI accessions via best-hit BLAST matching.

### Horizontal Gene Transfer (HGT) Identification

HGT predictions were computed using DarkHorse v2.0 (36,38,39) against the NCBI nr database (August 2024). Proteins were aligned using BLASTP (-evalue 1e-3, -max_target_seqs 100000) and self-matches were excluded using the -e option. DarkHorse was run with parameters -f 0.1 -n 3 -b 0 -d 1. Candidates were selected based on optimized lineage probability index (LPI) thresholds **(Supplemental Figure 1)** and manually verified. Functional annotations and taxonomic lineage assignments were generated using KEGG/BlastKOALA (40) and genes were grouped by function for downstream analysis.

### Bacterial Growth and Biodegradation Assays

Bacterial strains were grown overnight in Lysogeny Broth (LB) at 37°C. The five-member consortium was inoculated to a final **OD**_600_ of 0.01 into 1.5-cm glass tubes containing 5 mL of LCFBM medium in triplicate. The LCFBM base consisted of 10 mM sodium phosphate buffer (pH 7.4) supplemented with 0.05% (w/v) yeast extract, 0.2% (w/v) (NH_4_)_2_SO_4_, and 1% (vol/vol) trace elements (0.1% [wt/vol] FeSO_4_·7H_2_O, 0.1% [wt/vol] MgSO_4_·7H_2_O, 0.01% [wt/vol], CuSO_4_·5H_2_O, 0.01% [wt/vol] MnSO_4_·5H_2_O, and 0.01% [wt/vol] ZnSO_4_·7H_2_O). For PET plastic growth, cultures were grown in triplicate in LCFBM supplemented with 1% (wt/vol) of PET granules (Sigma-Aldrich, St. Louis, MO) and grown statically for 7 weeks; growth was monitored by CFU/mL counts. For investigation of MHET degradation, 5mL tubes containing LCFBM were supplemented with 0.2% (wt/vol) MHET (Advanced ChemBlock Inc, Hayward, CA) and incubated aerobically for 21 days.

### Proteome Extraction and Analysis

Extracellular proteins from culture supernatants and PET biofilms were extracted using high-salt ECM disruption (1 M NaCl) to avoid cell lysis. Culture supernatants (25 mL) were cleared (8,000 × g, 10 min, 25°C), adjusted to 1 M NaCl, and incubated for 15 min. Biofilm proteins were extracted from five PET pellets by vigorous washing in 1 mL of 1 M NaCl for 15 min. Both extracts were clarified at 5,000 × g for 10 min. Proteins were precipitated overnight at –20°C using four volumes of cold acetone. The resulting pellets were centrifuged (13,000–15,000 × g, 10 min), air-dried, resuspended in 100 µL ultrapure water, and stored at –20°C. Samples were subsequently analyzed by LC-MS/MS performed at the OHSU Proteomics Core Facility. Proteins were identified by searching against specific UniProt (41) reference proteomes, *Bacillus albus* (UP000181873), *B. thuringiensis* (UP000195991), and *Pseudomonas* spp. (UP001231507, UP000238367).

### Quantification of Aromatic Products (HPLC-UV)

Aromatic depolymerization products were quantified using an Agilent 1290 Infinity HPLC (Agilent Technologies, Santa Clara, CA) equipped with a diode array detector (DAD). Separation was performed on a ZORBAX Eclipse Plus C18, 100 × 2.1 mm, 1.8 µm column (Agilent Technologies, Santa Clara, CA) at 40°C. An isocratic mobile phase of water:acetonitrile (80:20) acidified with 0.1% (w/v) formic acid was applied at 0.450 mL/min, with UV detection at 250 nm. Both standards (TPA, MHET, BHET) and experimental samples were solubilized in 90:10 acetonitrile:DMSO prior to injection.

### Identification of Unknown Metabolites (LC-MS)

To identify non-standard compounds and intermediates, culture samples were diluted in 100% acetonitrile and analyzed using Shimadzu 8030 Triple Quadrupole LC/MS system (Shimadzu, Kyoto, Japan). Chromatographic separation was performed using the same mobile phase described above with an Agilent InfinityLab Poroshell 120 EC-C18, 4.6 x 100 mm, 4 µm column (Agilent Technologies, Santa Clara, CA). Mass spectral data were acquired in negative electrospray ionization (ESI) mode to structurally characterize aromatic byproducts and unknown intermediates generated during biodegradation. TPA and MHET standards ESI peaks were experimentally determined to be at 165.02 *m/z* and 209.05 *m/z*.

## RESULTS AND DISCUSSION

### Analysis of Horizontal Gene Transfer Events

The bacterial consortium possesses a large accessory genome (35) rich in diverse hydrolytic and oxidative enzymes involved in the biodegradation of plastics (28,42,43). As accessory genes are often transmissible, we hypothesized these functional traits may have been acquired from distant taxa (44). To test this hypothesis, we used DarkHorse(36) to detect atypical evolutionary signatures. The tool calculates a Lineage Probability Index (LPI), where a low score indicates that a gene’s closest homologs are found in distant taxa rather than the host lineage. This identifies probable HGT events based on atypical ancestry, regardless of transfer direction.

We identified and filtered putative HGT candidates **(File S1)**. Apparent eukaryotic hits were likely false positives caused by incorrect taxonomic annotation of contaminated WGS contigs (e.g., bacterial sequences labeled as eukaryotic), as evidenced by low similarity to true eukaryotes but strong similarity to bacterial query groups such as *Bacillus* or *Pseudomonas*. Approximately 13% of all predicted HGT candidates with LPI scores lower than 0.1 were phage-derived, predominantly within *Bacillus* strain 13.1 **(Table S1)**. After stringent filtering and removal of false positives and bacteriophage hits across the five strains, protein-coding density and predicted horizontal gene transfer (HGT) frequencies varied but remained within ranges typical of bacterial genomes **(Table 1).** These data show that up to 6.8% of genes within consortium members could be attributed to HGT alone and should be examined further to assess if they were implicated in PET depolymerization.

**TABLE 1.** NCBI genome statistics and summary of predicted HGT candidates’ genes from the five-member consortium.

| *Genome Statistics* | *Bacillus* Strains | *Pseudomonas* Strains |
| --- | --- | --- |

|  | 9.1 | 13.1 | 9.2 | 10/13.2 |
| --- | --- | --- | --- | --- |
| Total Protein CDS Genes | 5513 | 6577 | 5693 | 5220 |
| DarkHorse Predicted HGT Genes | 216 | 365 | 46 | 233 |
| DarkHorse % Predicted HGT | 4.1% | 6.8% | 0.8% | 4.5% |

Counts of total coding sequences (CDSs), initial DarkHorse HGT predictions of putative horizontally acquired genes, including phage, excluding false positives within each consortium member. 10/13.2, are 99.9% identical with minor strain differences.

HGT predicted genes in each strain were grouped by taxonomic distribution and abundance. Illustrated in **Figure 1**, HGT candidates were predominantly detected in Gammaproteobacteria for the *Bacillus* strains 9.1 and 13.1, with 197 and 266 genes, respectively, whereas HGT predicted genes in the *Pseudomonas* strains 10/13.2 (strains 10 and 13.2 show 99.9% DNA sequence identity) were predominantly predicted to be *Bacilli*-derived, with up to 143 genes detected in *Pseudomonas* strain 10. On the individual strain level, both *Bacillus* strains had highly similar HGT events (apart from multiple bacteriophage elements seen in strain 13.1 **(Figure S1)**). In contrast, *Pseudomonas* 9.2 strains differed dramatically from 10/13.2, with minimal detected HGT events of 0.8% (**Table 1)**. These results indicated that many genes were acquired via HGT from two distinct taxa: *Gammaproteobacteria* and *Bacilli*.

**Figure 1.**
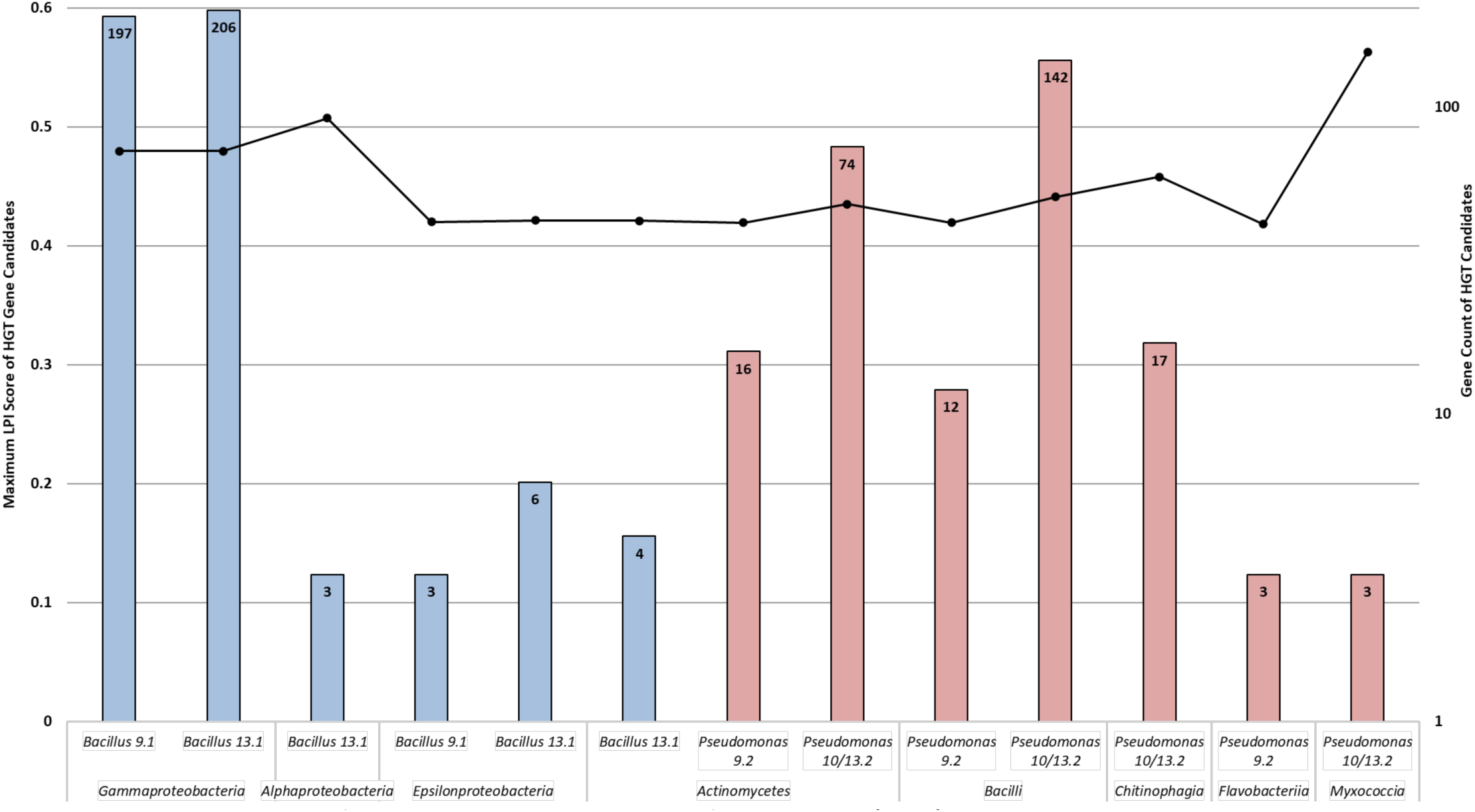
LPI distribution of predicted horizontally transferred genes (HGT) across consortium genomes. Gene counts (bars) and maximum LPI scores (line) for donors contributing > 2 genes (LPI cutoff < 0.6; viral/spurious hits removed). Blue: *Bacillus* strains (9.1, 13.1) show enrichment from *Gammaproteobacteria* (197 and 266 genes). Red: *Pseudomonas* strains (9.2, 10/13.2) show enrichment from *Bacilli* (12 and 142 genes), indicating bidirectional cross-class transfer.

Because DarkHorse donor assignments rely on global compositional similarity and because publicly available sequence databases are dominated by clinically relevant, not environmental taxa, predictions more frequently default to high-level matches at the phylum or class level rather than the genus or species level. As a result, DarkHorse may correctly identify donors/recipients at a higher taxonomic level, but genus and species assignments might be inaccurate. This led to secondary taxonomic and functional annotation of the unique HGT candidates using GhostKOALA(40).

Functional annotation was assigned to 45% of HGT candidates. The genes were annotated as having core metabolic and housekeeping functions **(Figure 2)**. Because single-gene predictions can reflect spurious annotations or transient genetic acquisitions, we prioritized genes from taxa that were detected multiple times for further analysis. Recurrence suggests stable inheritance or repeated acquisition events, and therefore a higher likelihood of biological relevance than single-copy occurrences. Most HGT genes were implicated in cellular processes **(Figure 2)** and were not assigned to xenobiotic metabolism. Functional and taxonomic classifications indicated high-frequency interspecies transfer between consortium members.

**Figure 2.**
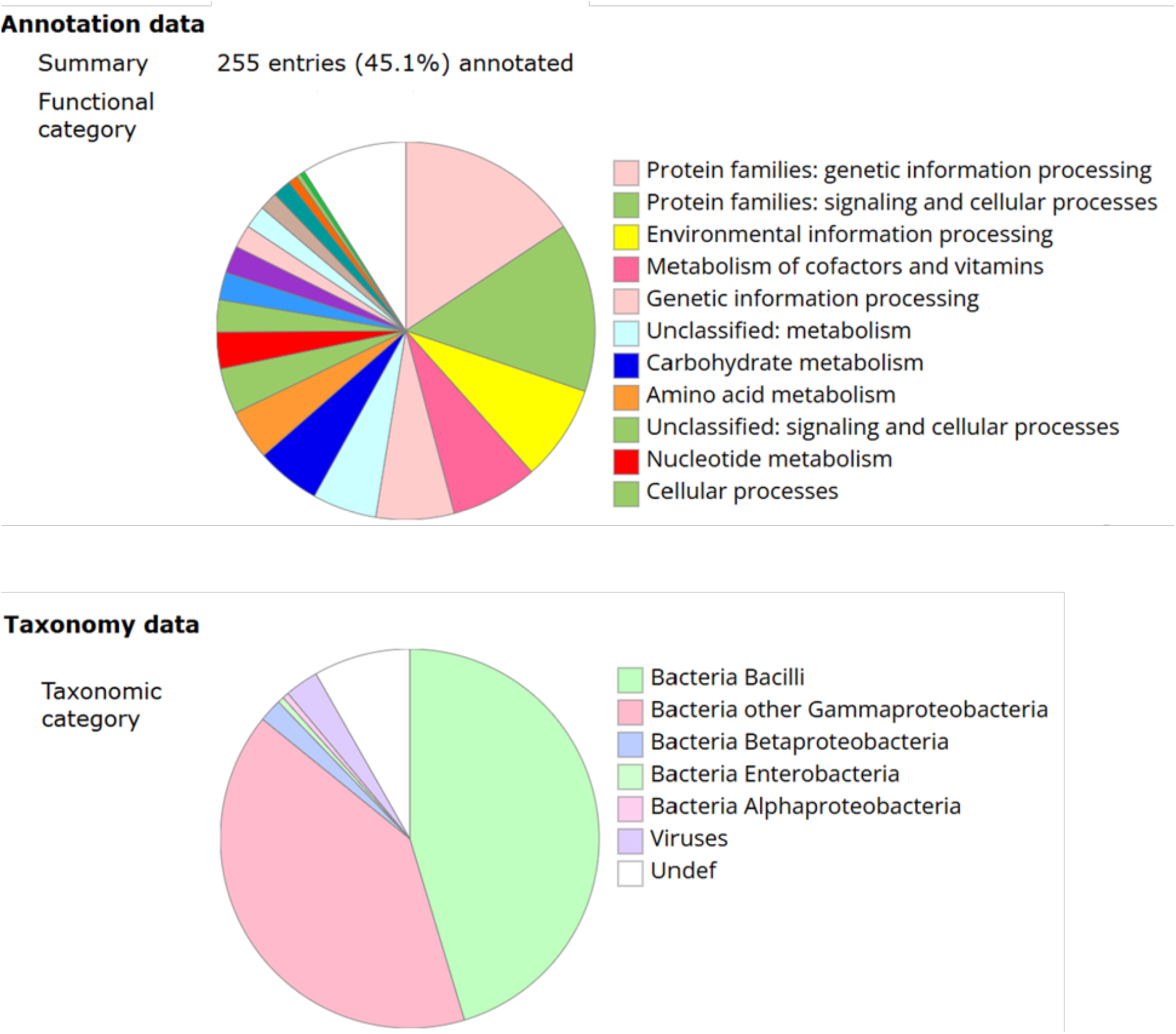
Functional and taxonomic annotation of HGT candidates using GhostKOALA. Functional annotations of all unique HGT genes (top) were assigned KEGG-based pathway assignments, of which 255 genes (45.1%) received functional annotations. The top categories were comprised of genetic information processing, signaling/cellular processes and environmental information and processing. Taxonomic annotation (bottom) shows the predicted lineage of the closest homologs, with the majority assigned to Bacilli and Gammaproteobacteria, with some smaller contributions from α-, β-, and Enterobacteria. Minor viral and undefined assignments were also observed. (KEGG Figure)

The majority (>85%) of HGT candidates were classified as either *Bacillus* or *Pseudomonas* in origin **(File S2)**. Minor contributions from other taxonomic lineages were observed **(Figure S2),** including seven genes from non-*Pseudomonas Gammaproteobacteria* (*Acinetobacter, Aliivibrio, Azotobacter, Shewanella, Vibrio*) and five genes from non-*Bacillus Bacilli* (*Lacticaseibacillus, Listeria, Staphylococcus*). Functional annotation revealed that these specific transfers encoded mostly genetic information for processing proteins (*e.g.,* peptide deformylase, S1 ribosomal protein) and hypothetical proteins rather than key metabolic enzymes involved in PET biodegradation. The metabolic exception was a *pcaD* gene (TPA metabolism) (45)) from *Azotobacter*. These data indicate that HGT reflects localized gene exchange rather than a primary mechanism for evolving hydrocarbon degradation pathways. Shown in **Table 2** are the comparisons between DarkHorse predictions^±^ and KEGG lineage predictions˟ between the queried genes within the bacterial consortium^+^. Functional enzymes* within this pangenome include hydrolases, oxidoreductases, and associated regulators and transporters **(Table S2)**. (35).

**TABLE 2.** Taxonomic and functional classification of horizontally transferred accessory genes within the consortium.

|  |  |  |  |  |
| --- | --- | --- | --- | --- |
| Query Genus<sup>+</sup> | DarkHorse lineage<sup>±</sup> | KEGG lineage<sup>×</sup> | Functional category<sup>*</sup> | Gene annotation examples |
| *Pseudomonas* | Bacilli | γ-proteobacteria | Glycosylation | GT-A fold GT, PimA-like |
| *Pseudomonas* | γ-proteobacteria | γ-proteobacteria | Hydrolases | PcaD (3-oxoadipate enol-lactonase) |
| *Pseudomonas* | γ-proteobacteria | γ-proteobacteria | Oxidoreductases | α-ketoglutaric semialdehyde dehydrogenase, NirD |
| *Bacillus* | γ-proteobacteria | γ-proteobacteria | Oxidoreductases | Cytochrome P450 |
| *Pseudomonas* | Bacilli | γ-proteobacteria | Oxidoreductases | Phthalate dioxygenase reductase-like, SDR enzymes |
| *Bacillus* | γ-proteobacteria | Bacilli | Oxidoreductases | NADH dehydrogenase, QueG |
| *Pseudomonas* | Bacilli | γ-proteobacteria | Regulators | SutA, LysR-family transcriptional regulators |
| *Bacillus* | γ-proteobacteria | Bacilli | Regulators/Structural | CstA carbon starvation protein, SpaA pilin-related |
| *Pseudomonas* | Chitinophagia | β-proteobacteria | Transporters | CrgA-like PBP2 |
| *Pseudomonas* | Bacilli | γ-proteobacteria | Transporters | TRAP transporter, periplasmic binding proteins |

Accessory genes from consortium members implicated in plastic biodegradation were compared with taxonomic assignments derived from DarkHorse and KEGG-based annotations to evaluate consistency and potential donor misassignment. Genes were summarized by query genome<sup>+</sup>, DarkHorse predicted donor class<sup>±</sup>, and KEGG-inferred lineage<sup>×</sup>, then grouped by functional category<sup>\*</sup>. Functional categories highlight enrichment in hydrolases, oxidoreductases, transporters, and regulatory proteins associated with aromatic PET-monomer metabolism.

Finally, because many genes may contribute indirectly to plastic biodegradation by supporting surface attachment, stress tolerance, and monomer assimilation, we examined HGT candidates associated with these processes. In *Bacillus* strains, horizontally transferred genes related to biofilm formation (*spaA*), stress tolerance, and starvation responses (*cstA*) were identified, traits that increase adhesion to polymers and enhance persistence under nutrient limitation (46–48).

In *Pseudomonas* strains, several oxidoreductases and transporters associated with aromatic compound metabolism were detected as HGT candidates, including *pcaD*, as well as dehydrogenases and transcriptional regulators potentially involved in terephthalate and ethylene glycol metabolism (49–54). Homologs of some of these genes were also detected in *Actinomycetota*, including *Arthrobacter*, taxa commonly associated with petroleum-contaminated environments. However, KEGG-based classification suggests these loci are more likely native to Gammaproteobacteria (52,55). Importantly, no complete gene cassettes associated with PET depolymerization were identified via DarkHorse analysis; instead, HGT appears to contribute individual genes that enhance existing metabolic flexibility, stress resilience, and surface-associated functions, rather than acting as a primary mechanism for disseminating canonical plastic-degrading enzymes such as PETases.

### Mobile Genetic Elements Suggest Xenobiotic Specialization

Most predicted HGT events occurred within the same genus (e.g., *Pseudomonas* and *Azotobacter*) or between closely related taxa, consistent with short-range gene flow within biofilms (46,56,57). Although some distantly acquired genes were detected, the overall pattern supports lineage-specific refinement rather than widespread long-distance transfer (58,59). Because xenobiotic pathways, including TPA metabolism, in *Pseudomonas* are often plasmid-borne (60–63), we screened all strains for plasmids and other mobile genetic elements using IMG/JGI and geNomad. Additionally, as regulatory and metabolic functions are typically associated with larger genetic cassettes, we examined all predicted plasmids, transposable elements, and phage-associated genes using genome annotations and via geNomad (64), an AI tool that classifies MGE through the IMG/JGI portal (65).

Among *Bacillus*, strain 9.1 contained a small 5.6 kb plasmid encoding a toxin–antitoxin module, while strain 13.1 harbored phage-associated elements but no large accessory catabolic plasmids. *Pseudomonas* strain 9.2 lacked predicted plasmids but contained scattered transposases. In contrast, *Pseudomonas* strains 10/13.2 shared a large 247,856 bp genomic island **(File S3)** with high similarity to plasmids from other *Pseudomonas* spp., including *Pseudomonas putida S12* (0.834), a solvent-tolerant strain that can degrade styrene plastics (62,66,67). This genomic island demonstrates that strain 10/13.2 engaged in direct transfer between species within the same genus. Given its size and presence of plasmid stabilization proteins (e.g., ParE) and catabolically relevant genes, including hydrolases, aryl-alcohol dehydrogenases, oxidoreductases, and redox enzymes, this island was examined for TPA metabolic potential (20,45).

Shown in **Figure 3**, complete benzoate and phenylacetate degradation cassettes were present, enzymes associated with phenylpropanoid and phthalate metabolism, and multiple aromatic transporters, including a plasmid-associated *pcaK* homolog previously implicated in TPA uptake(45,68,69). The coexistence of ABC-type uptake systems and MFS-family efflux transporters indicate regulated aromatic handling. Stress-response genes (*lexA*, *imuA*) are also present, potentially enhancing tolerance to xenobiotic and solvent stress. The presence of both ABC-type uptake systems and multidrug/MFS-family efflux transporters suggests bidirectional trafficking of aromatic substrates or metabolites.

**Figure 3.**
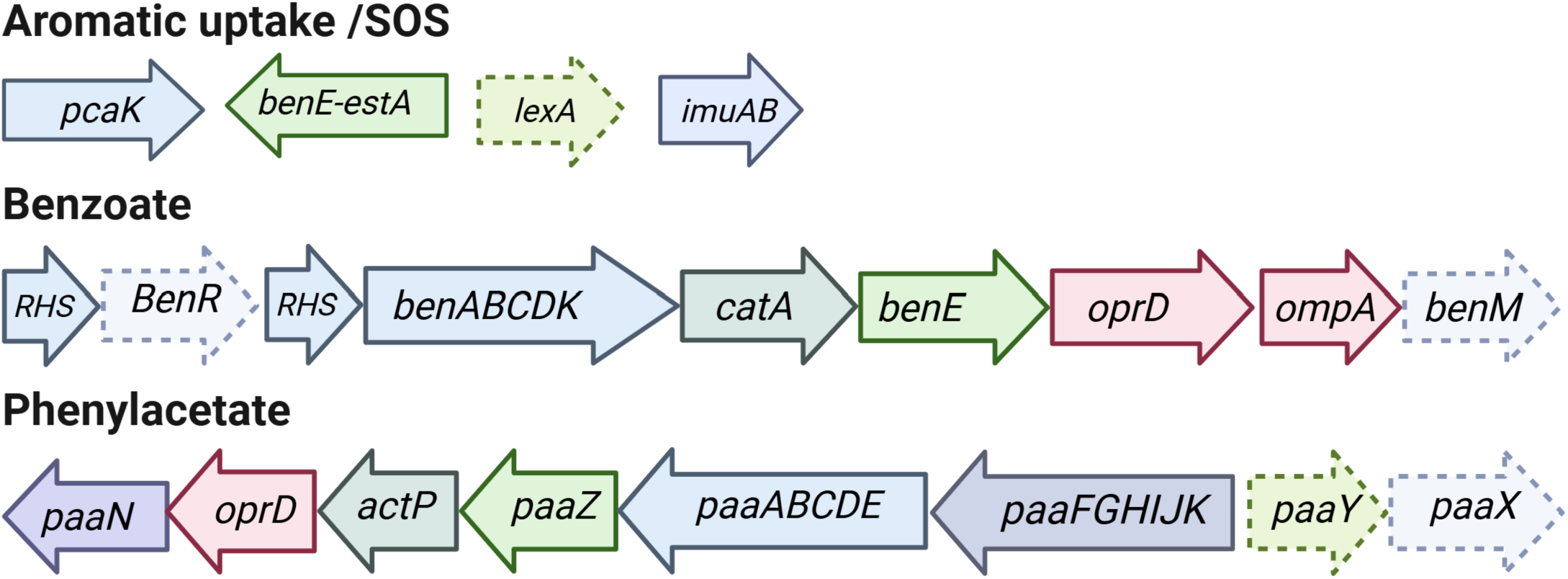
Functional gene cassettes within xenobiotic-associated genomic island. Gene clusters are linked to aromatic metabolism, stress tolerance, and plasmid maintenance. A 4-hydroxybenozate-like (*pcaK*) transporter was found encoded next to SOS (*lexA–imuA–imuB*) genes. Catabolic modules were annotated as benzoate/catechol degradation (*benABCD–catA–benE*), and phenylacetic acid catabolism (*paaZ-paaABCDE-paaFGHIJK-paaY-paaX*). Arrows indicate gene orientation. Dashed arrows indicate regulatory elements. Genomic regions shown in **Table S3** in *Pseudomonas* strain 10.

Several additional conserved but poorly characterized proteins (DUF1302, DUF1329) are interspersed within the cassette, which have been implicated in fatty acid metabolism, potentially containing esterase-like domains. Individual genes show homology to components of phenylpropionate, flavonoid (e.g., naringenin), lignin-derived aromatic, and phthalate-associated pathways. However, the overall organization is not typical of aromatic degradation modules, normally having more genetic organization. **File S3).** These data suggest that this cassette functions as a broad-substrate aromatic catabolism system. Notably, *Pseudomonas* strain 9.2 lacks this island yet retains PET-degrading capability, indicating that core chromosomal pathways within the *Pseudomonas* strains are sufficient for basal PET breakdown. The plasmid-derived island in strains 10/13.2 appears to represent a secondary adaptive acquisition that enhances aromatic assimilation efficiency, including TPA handling. Although originally mobilized via horizontal transfer, the locus now appears genomically stabilized. These findings point towards an evolutionary strategy where accessory modules for aromatic handling act alongside core degradation pathways, which likely increases metabolic synergy within the consortium.

### Proteomic and Functional Validation

To validate prior multi-omics predictions and to identify key enzymes mediating PET depolymerization and catabolism of monomers, we used proteomics to profile proteins actively expressed and secreted within PET-biofilms of the full consortium (FC). We examined the proteome from the media, as well as proteins extracted from the extracellular matrix of the biofilms found on the surface of PET after incubation. We saw distinct proteomic signatures from the different strains, consistent with the observed synergy **(Files S4 and S5).**

*Bacillus* are well known as prolific protein secretors, biofilm producers, and biosynthetic specialists, particularly in the production of extracellular enzymes and secondary metabolites (120–122). Consistent with these traits, the fraction of *Bacillus*-derived proteins assigned to general metabolic pathways was primarily associated with biosynthesis of secondary metabolites and metabolism of cofactors and vitamins **(Figures 4A and S3A).** Beyond core metabolic contributions, *Bacillus* strains accounted for the largest proportion of enzymes not assigned to central metabolic pathways and were enriched in proteins involved in protein modification. The *Bacillus* proteome was dominated by secreted hydrolases, oxidoreductases, and proteolytic enzymes, supporting a functional role in extracellular polymer and oligomer modification at the PET surface rather than in downstream metabolic assimilation **(Figures 4B and S3B).** In addition to these catalytic proteins, we observed specific biofilm and surface proteins, including CalY biofilm matrix protein and SpaA pilin-related proteins from the *Bacillus* strains **(Table 3).**

**Figure 4.**
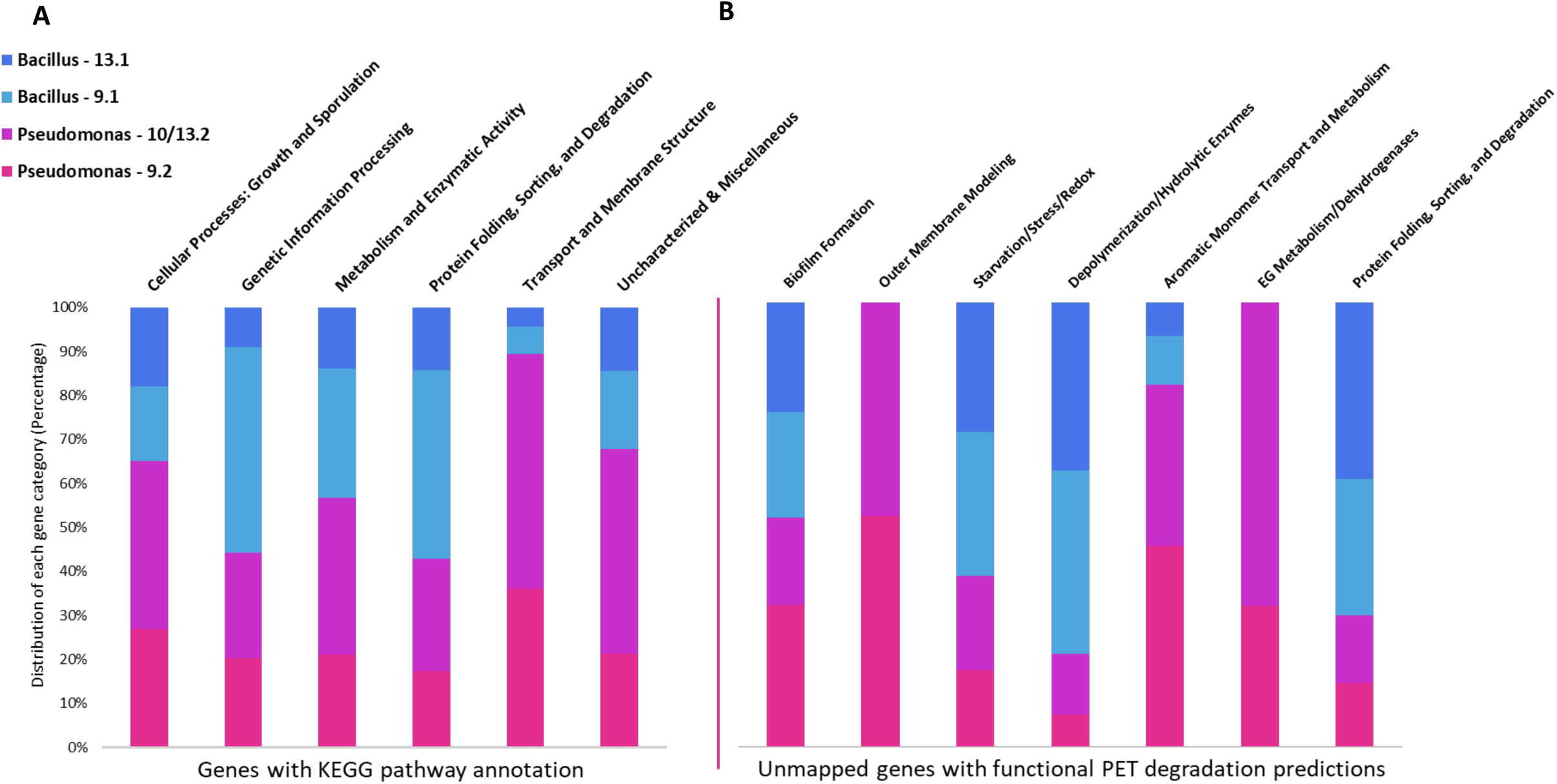
Proteome metabolic pathway mapping and functional annotation of PET-relevant categories between consortium members show division of labor when grown on PET. Functional roles of consortium members were examined using proteome functional annotation. KEGG Orthology (KO) assignments and general protein functions were mapped to KEGG pathways and consolidated into 6 general categories **(File S7)**, cellular processes: growth and sporulation, genetic information processing, metabolism and enzymatic activity, protein folding, sorting and degradation, transport and membrane structure, uncharacterized & miscellaneous **(A)** uncharacterized & miscellaneous proteins, and proteins unassigned KO, were sorted and functional predictions were assigned into broad categories based on gene annotation and function within each strain **(B).** Categories include biofilm formation, outer membrane modeling, starvation/stress/redox, depolymerization/hydrolytic enzymes, aromatic monomer transport and metabolism, EG metabolism/dehydrogenases and protein folding, sorting, degradation **(File S8).** Undefined genes are not shown. Bars represent total proteins from each strain and the proportion they contribute overall, highlighting strain-specific function.

**Table 3.** Non-core annotated proteins with associated plastic colonization and biodegradation functions.

| Functional Category<sup>‡</sup> | *Pseudomonas* | *Bacillus* |
| --- | --- | --- |
| **Nitrogen Metabolism**<sup>(70–73)</sup> | Aminotransferases (DavT, AspAT);<br>Glutamine amidotransferase;<br>Carbamoyltransferase; Nucleobase<br>deaminase | Aminopeptidases (M18, M42,<br>Xaa-Pro) |
| **Biofilm & Quorum Sensing**<sup>(74–83)</sup> | AHL acylase; Adhesins; Polysaccharide<br>export; PHA granule protein; Iron OM<br>receptors | Adhesins (SpaA); Collagen-<br>binding protein; Camelysin |
| **Ethylene Glycol Metabolism**<sup>(84–89)</sup> | Aldehyde & alcohol dehydrogenases;<br>Quinohemoprotein dehydrogenase;<br>Acyl-CoA dehydrogenase; PQQ synthase | — |
| **Genetic Information Processing** | Topoisomerase II; NusA; IF-2/IF-3; Fur;<br>H-NS | DNA polymerase X; HU;<br>Exonuclease; IF-3 |
| **Growth & Regulation**<sup>(47,90)</sup> | MreB; MinD; P-II regulator; Elongation<br>factors | Sporulation proteins (SpoVG,<br>SpoE, GerQ); FtsN |
| **Hydrolytic / Oxidative Enzymes**<sup>(91–99)</sup> | Chloroperoxidase; Dienelactone<br>hydrolase; Hydrolases | Chloroperoxidase;<br>Nitroreductase; Azoreductase;<br>Endoglucanase; Metal-<br>dependent hydrolases |
| **Outermembrane Transport /**<br>**Aromatic Transport**<sup>(100–107)</sup> | OprD/Omp porins; MFS & ABC<br>transporters; TolQ/TolB; MlaA; PqiB; OM<br>esterase (EstP); FA transporters | Iron/heme transporters;<br>Siderophore systems; Cation<br>symporters |
| **Protein Folding / Degradation**<sup>(108–110)</sup> | HslUV protease; Trigger factor;<br>Oligopeptidase; Membrane dipeptidase | PrsA foldase; Serine protease;<br>Aminopeptidases;<br>Oligoendopeptidase |
| **Redox Handling**<sup>(111–115)</sup> | Azurin; Cytochromes; Copper oxidase;<br>ETF; RidA; Carbonic anhydrase | Cytochrome c oxidase;<br>Peroxiredoxin |
| **Stress & Starvation**<sup>(116–119)</sup> | CstA; Dps-like DNA-binding protein;<br>EttA; Response regulators | Dps-like protein; Acid<br>phosphatase |
| **Undefined** | DUF families (1302, 1329, 3108, etc.);<br>Hypothetical; Phage-associated | DUF2642; DUF1641;<br>Hypothetical |

<sup>a</sup>Functional categories were identified from relevant literature cited; **File S6** contains a comprehensive list of all non-core proteins found within the proteome annotated by KEGG.

In contrast, in the *Pseudomonas* strains, proteome abundance included numerous porins and transporters (Multiple OprD family, ABC and TolC/RND Tefflux shown in **Table 3**), indicating active transport of compounds such as TPA (106). The expression of multiple MFS transporters facilitates active import of aromatic compounds (123,124) **(Table 3)**. Expression of SAM-dependent methyltransferase and hydrolases, including previously identified EstP (35) are also present and are indicative of methyltransferase and hydrolysis activity, which may be directly involved in the hydrolysis of PET oligomers. Abundant alcohol and aldehyde dehydrogenases show active detoxification and metabolism of alcohols. This indicates that EG metabolism is only shown to be active in *Pseudomonas* strains **(Figure 4B).** Overall, this proteomic data shows that *Bacillus* are actively involved in the colonization and hydrolysis of the PET polymer and BHET oligomer, while *Pseudomonas* play an important role in the breakdown, detoxification, and assimilation of oligomers (*i.e.*, MHET) and monomers (TPA and EG).

### MHET modification expands PET oligomer assimilation pathways

Previously, ¹H NMR analysis of PET leachates following consortium incubation revealed unique methyl, glycol, acetaldehyde, and carboxylic acid resonances absent from abiotic controls, indicating operation of a specialized enzymatic cascade that couples PET oligomer hydrolysis with downstream modification to produce small, metabolizable aldehydes and organic acids. Interestingly, genome- and proteome-guided analyses did not identify a clear canonical MHETase homolog across consortium members, raising the possibility that MHET represents a potential metabolic bottleneck. Accordingly, we incubated individual isolates and the full consortium with MHET as the sole added substrate to assess whether MHET is directly hydrolyzed, transformed through alternative routes, or preferentially processed by specific community members.

Strain-specific MHET degradation and transformation were observed over the course of 21 days. When all consortium members were incubated with MHET as a carbon source, nearly complete hydrolysis of MHET was achieved by the full consortium, but not individual strains, showing synergistic degradation and mirroring those of PET and BHET as observed previously(14,35,37). Most interestingly, we observed a secondary aromatic product in addition to the canonical TPA byproduct of MHET that was not observed in the abiotic control **(Figure 5, Figures S4 and S5).** Using LC-MS, determining mass, retention time, and fragmentation patterns, an ESI peak at 223.05 m/z indicates that this product is the methylated aromatic ester of MHET, methyl mono-(2-hydroxyethyl) terephthalate (MMHET) **(Table S4).**

**Figure 5.**
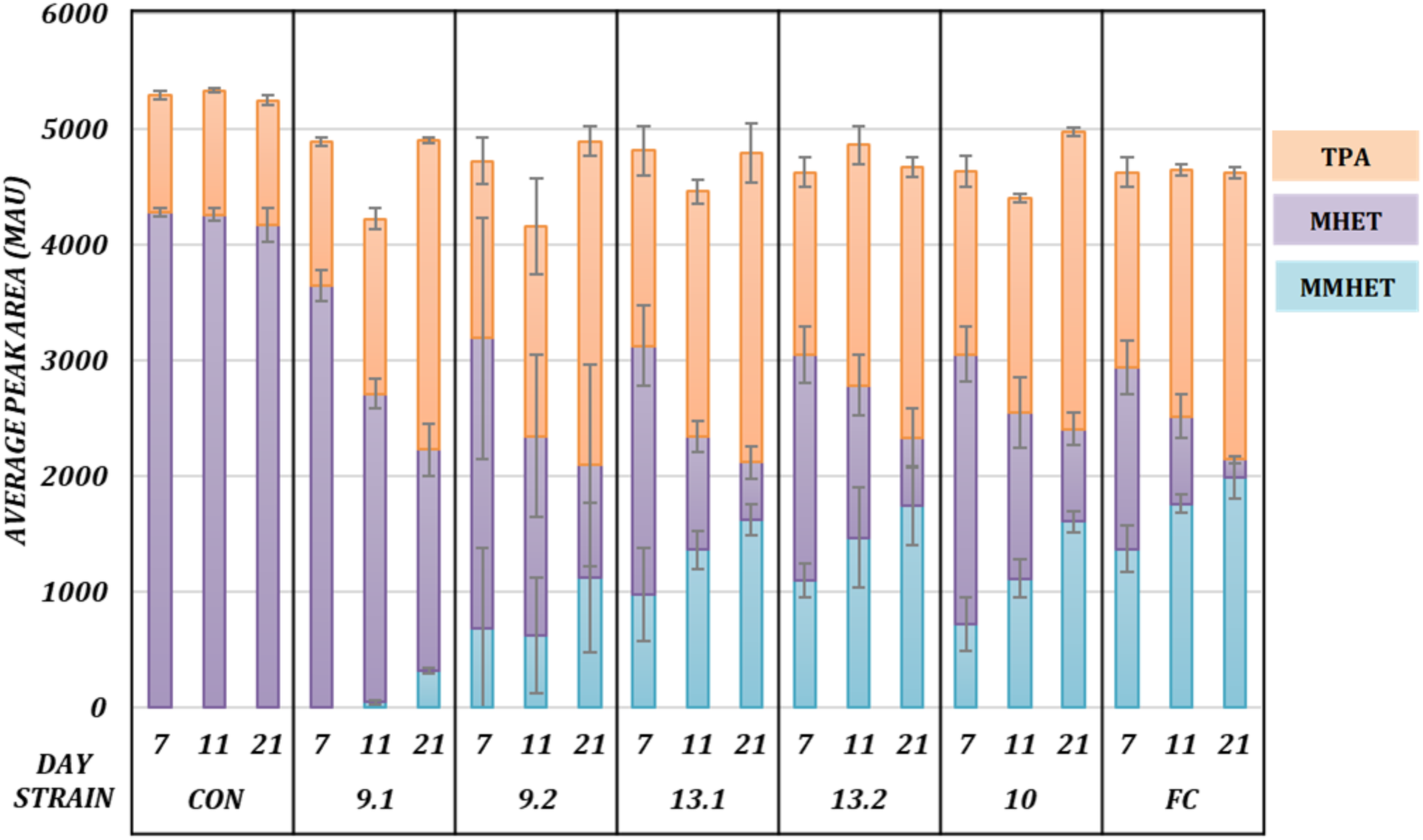
Quantification of MHET hydrolysis by HPLC indicates the full consortium (FC) synergistically degrades MHET. Hydrolysis of 0.2% (w/v) MHET was monitored over 21 days in liquid carbon-free basal medium (LCFBM) supplemented with 0.05% yeast extract. Variable residual MHET levels were detected across individual isolates, whereas depletion was only observed by day 21 in the Full Consortium (FC) containing all five strains, indicating that conversion of MHET to terephthalic acid (TPA) requires cooperative activity among consortium members. Control (CON) group quantifies spontaneous hydrolysis of MHET under culturing conditions over time. Error bars denote standard deviation (n=3).

These data suggested that when grown on the oligomer MHET, the strains methylate and hydrolyze MHET to reduce redox stress. MHET is not only an enzymatic inhibitor of PETase activity, but also an amphipathic aromatic ester that causes significant membrane stress and redox pressure. Thus, methylation, hydroxylation and carboxylation are typical strategies employed by HCB to degrade aromatic compounds, particularly within anaerobic biofilms. These findings show a modified enzymatic approach to PET degradation beyond hydrolysis, emphasizing methylation as a viable method for prolonged degradation. Products detected from the FC grown on PET are shown in **Figure 6** with the predicted enzymatic cascades identified from proteomic and genomic annotations.

**Figure 6.**
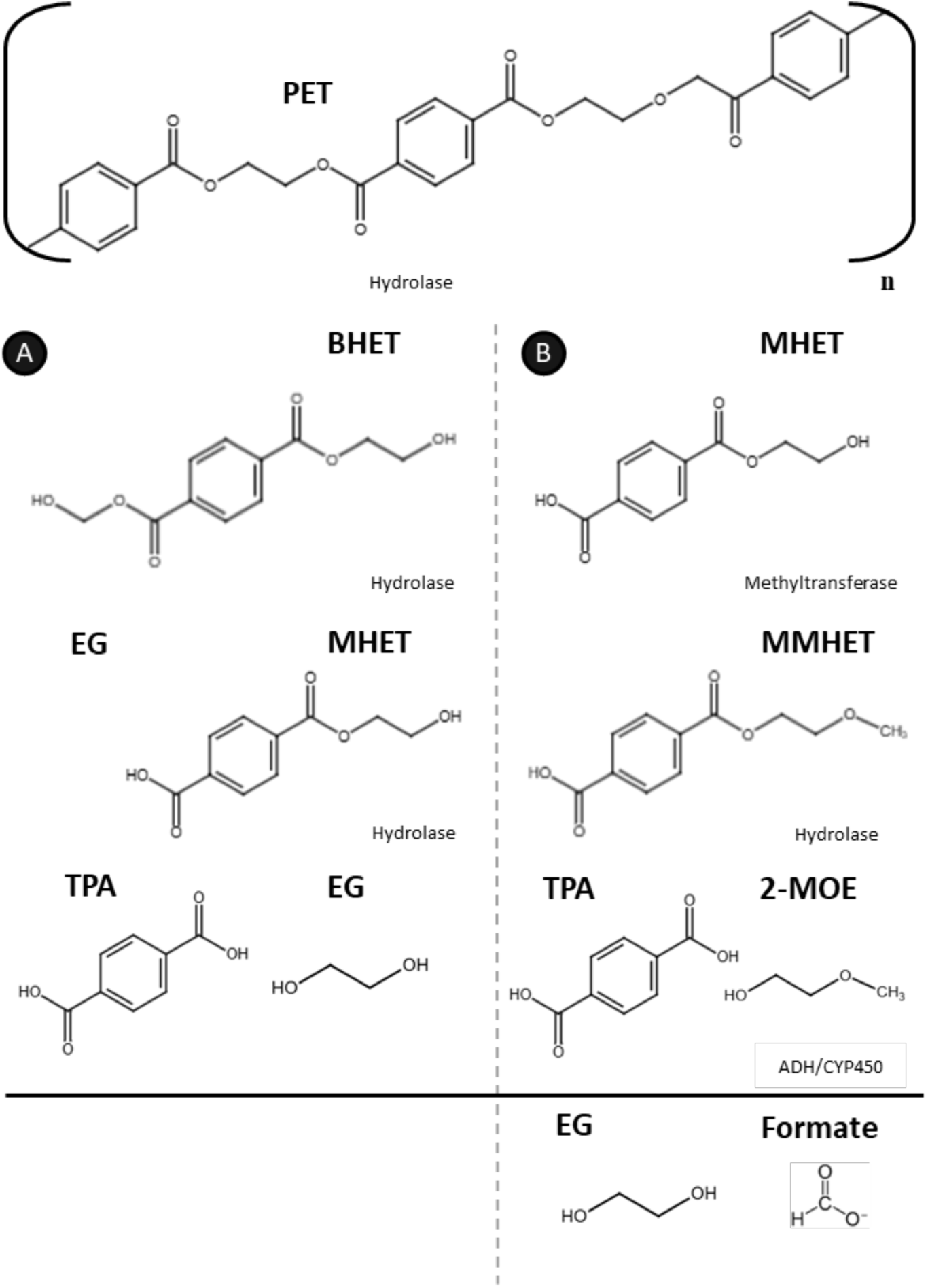
Proposed metabolic pathways for PET depolymerization and monomer assimilation by the full consortium. Schematic illustrating two complementary routes by which consortium members convert polyethylene terephthalate (PET) into metabolizable substrates. **(A)** In the canonical hydrolytic pathway, PET is depolymerized into the soluble oligomers is (2-hydroxyethyl) terephthalate (BHET) and mono(2-hydroxyethyl) terephthalate (MHET), which are further hydrolyzed to terephthalic acid (TPA) and ethylene glycol (EG). TPA is assimilated primarily by *Pseudomonas* strains, while EG can be utilized by multiple consortium members. **(B)** An alternative transformation pathway involves modified or methylated MHET derivatives (e.g., MMHET), yielding downstream products such as 2-methoxyethanol (2-MOE) and formaldehyde, which can enter central metabolism or detoxification pathways.

### Conclusions

Our findings indicate that polyethylene terephthalate (PET) biodegradation within the full consortium is an emergent, cooperative process structured around metabolic division of labor. Within this system, *Bacillus* strains primarily contribute extracellular hydrolytic capacity and excel at persistence under environmental stress, whereas *Pseudomonas* strains specialize in downstream aromatic assimilation and redox management. Neither genus alone achieves complete PET mineralization, underscoring that polymer degradation and assimilation in this system are inseparable from interspecies cooperation. Perhaps consistently, the *Bacillus* and *Pseudomonas* strains from petroleum-polluted soils were difficult to isolate, away from each other, into pure cultures when initially enriching for esterase activity (74). Such metabolic interdependence is consistent with the growing body of evidence that environmental plastic degradation is rarely mediated by single organisms, enzymes, or linear pathways, but instead arises from coordinated activity across microbial communities.

This cooperative strategy contrasts with previously characterized PET-degrading systems, with single or dual enzyme systems that characterize enzymatic, not the microbial-mediated or complex enzymatic cascades that enable the prolonged breakdown of PET in the environment. In contrast, the FC reflects an exaptive trajectory where enzymes and pathways originally evolved for hydrocarbon and ester metabolism appear to have been repurposed for plastic-associated substrates. Rather than evolving PET-specific enzymes de novo, consortium members appear to exploit pre-existing catalytic promiscuity, reinforced by ecological interactions and biofilm-associated lifestyles. This distinction highlights that plastic biodegradation can arise through multiple evolutionary routes depending on environmental history and selective pressures.

Genomic analysis supports a model in which short-range horizontal gene transfer (HGT) contributes to functional complementarity rather than wholesale pathway acquisition. Although many predicted HGT events were phage-associated (consistent with broader patterns of bacterial genome evolution), accessory genes implicated in aromatic metabolism, transport, regulation, and stress response were enriched within the consortium and frequently shared between *Bacillus* and *Pseudomonas*. Notably, a large xenobiotic-associated genetic island identified in two *Pseudomonas strains* encode extensive catabolic, transport, and regulatory modules linked to aromatic metabolism and stress tolerance. The size and composition of this island suggest it functions as a modular platform for metabolic flexibility, enabling rapid adaptation to aromatic xenobiotics such as the oligomers and monomers generated from PET depolymerization. However, the ability to degrade PET oligomer does not reside solely within this plasmid, showing increased consortium flexibility. Together, these observations indicate that HGT in this system acts to fine-tune existing metabolic networks rather than introduce entirely novel pathways.

Proteomic analysis under PET conditions reinforces a division of labor at the functional level. *Bacillus* strains are enriched in secreted hydrolases, peroxidases, and biofilm-associated proteins, consistent with roles in surface colonization and oligomer hydrolysis. In contrast, *Pseudomonas* strains exhibit enrichment of transporters, oxidoreductases, and stress-response enzymes, positioning them as primary hubs for aromatic assimilation and redox balancing. This asymmetry suggests that chemical constraints imposed by PET depolymerization, particularly the accumulation of acidic or redox-active intermediates, necessitate metabolic handoff between taxa to maintain a viable environment sustaining plastic depolymerization over time.

One such constraint is the handling of mono(2-hydroxyethyl) terephthalate (MHET), a key PET oligomer intermediate. In addition to canonical hydrolytic processing, our data support the existence of an auxiliary transformation route involving MHET methylation and associated redox chemistry, yielding methylated MHET (MMHET) and downstream alcohol or aldehyde intermediates. We propose that this modification functions as a detoxification and solubilization strategy, mitigating acidification and facilitating transport or further processing within the biofilm. Such chemical flexibility is consistent with the enrichment of SAM-dependent methyltransferases, oxidoreductases, and aldehyde dehydrogenases observed in the accessory genomes and proteomes, particularly within *Pseudomonas*. By distributing hydrolysis, detoxification, and redox management across community members, the consortium can accommodate fluctuating chemical conditions that would likely inhibit individual strains.

Rather than proceeding through a single linear pathway, PET biodegradation in this system appears to operate through branched, condition-dependent routes that balance hydrolysis, detoxification, and aromatic assimilation across taxa. This strategy reflects the exaptation of ancestral hydrocarbon metabolism within a cooperative ecological framework, allowing the consortium to tolerate chemical stress while promoting cross-feeding and metabolic specialization. Complete PET mineralization occurs only under consortium conditions, with accumulation of MHET when key members are absent, demonstrating that plastic degradation here is fundamentally a community-level trait. Together, these findings illustrate how microbial cooperation, evolutionary history, and chemical constraints intersect to enable adaptation to novel anthropogenic substrates in hydrocarbon-rich environments.

## ACKNOWLEDGEMENTS

The authors thank the OHSU Proteomics Core Facility for LC-MS/MS data and Libby Brennan for preparation of the bacterial cultures necessary for this analysis.

## DATA AVAILABILITY STATEMENT

The data are contained within the article.

## AUTHOR CONTRIBUTIONS

Conceptualization: J.L.M, S.E, and I.N. Investigation: S.E., D.R., and P.P.; Writing—original draft preparation: S.E. and J.L.M.; Funding acquisition: J.L.M and P.P. All authors have read and agreed to the published version of the manuscript.

