## Supplemental Figures and Tables for "Ancestral Hydrocarbon Metabolism Enables PET Degradation by a Natural Bacterial Consortium"

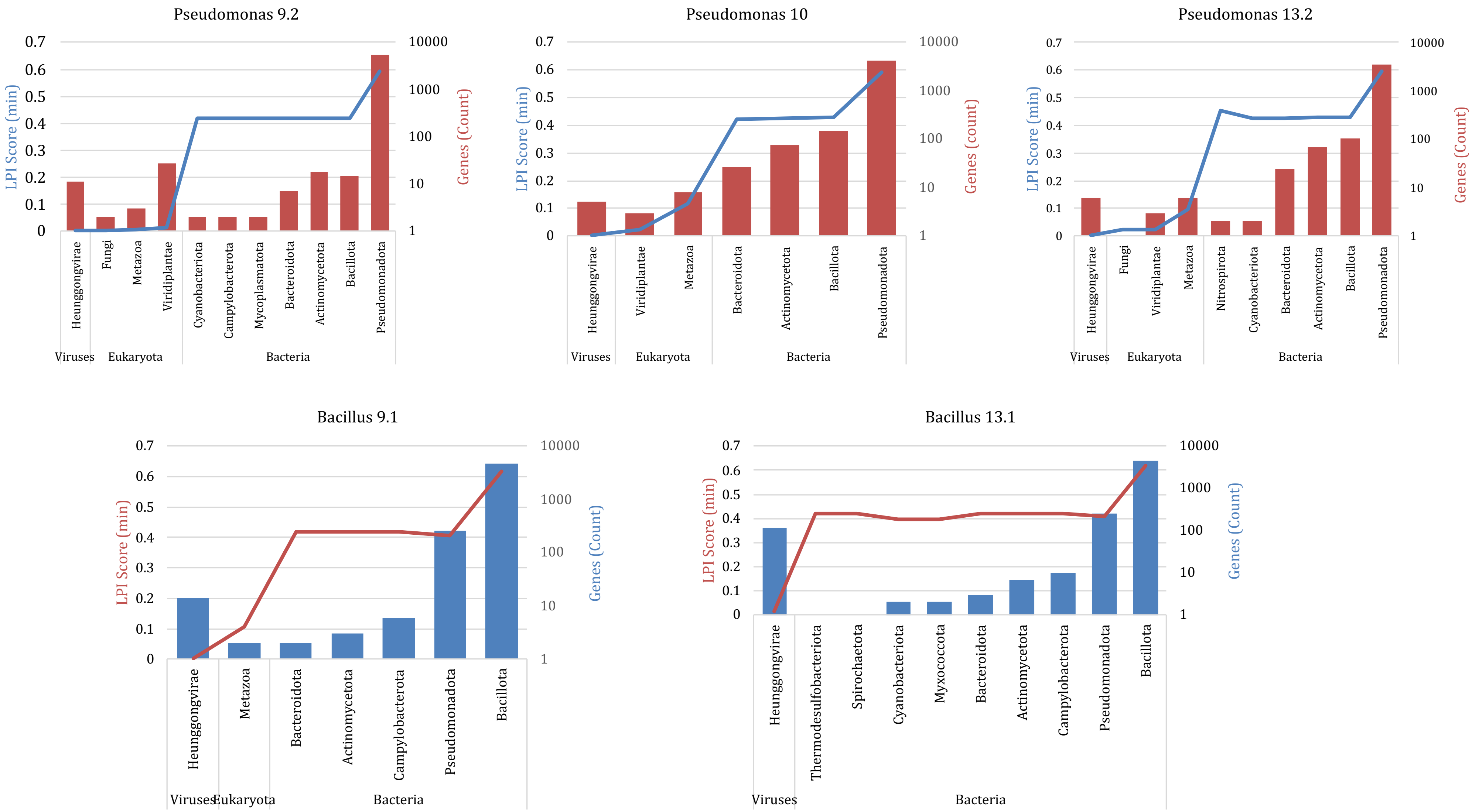


**Figure S1. Distribution of DarkHorse lineage probability index (LPI) scores across consortium member genomes.** For each consortium member, all predicted genes were assigned a lineage probability index (LPI) score using DarkHorse to assess potential horizontal gene transfer. Distributions show the frequency of genes across LPI values for each genome. Empirically determined thresholds of LPI ≤ 0.6 (broad cutoff) with <20% BLAST hits originated from self-identified taxa and LPI ≤ 0.4 (stringent cutoff) were used to identify candidate horizontally transferred genes. Lower LPI scores indicate reduced lineage consistency and a higher likelihood of horizontal acquisition.

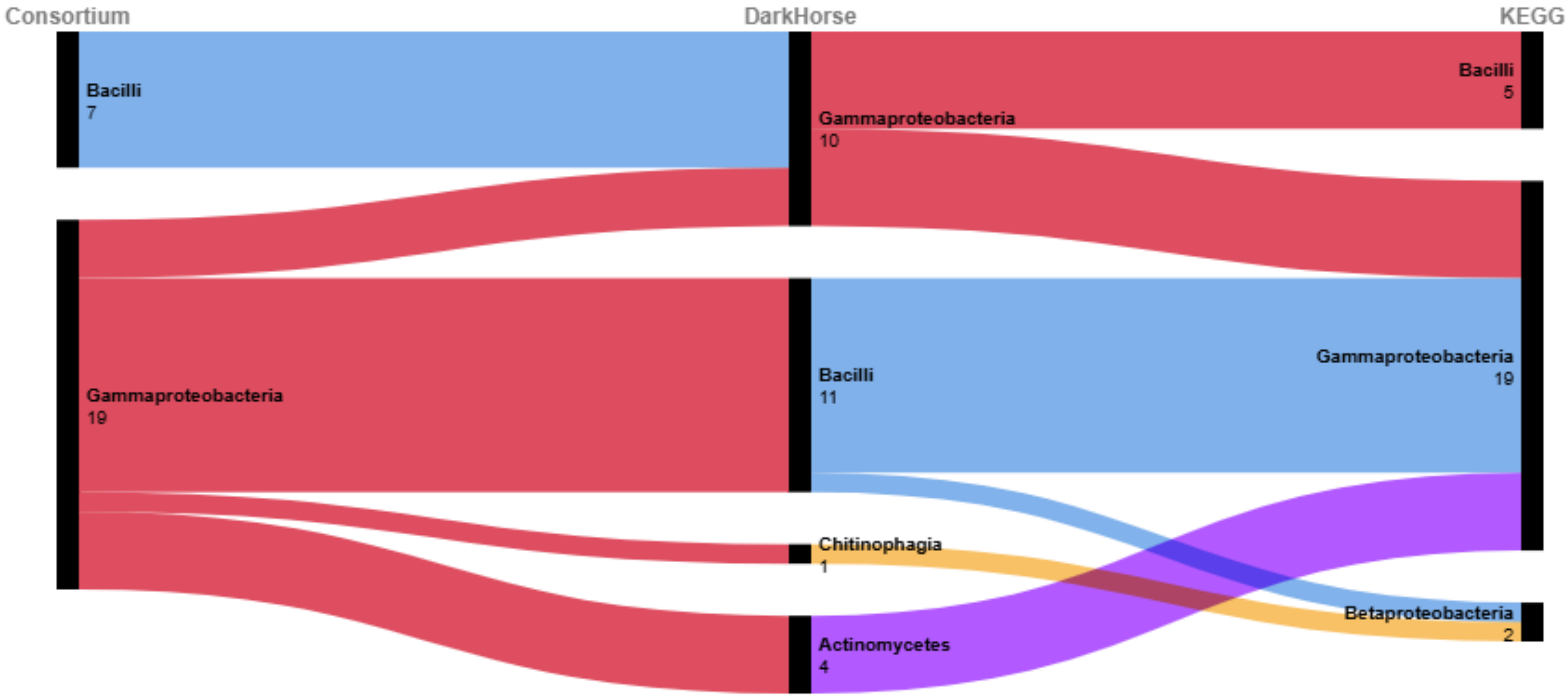


**Figure S2.** Sankey diagram illustrating relationships between consortium member taxa (left), DarkHorse-predicted donor lineages (center), and KEGG-based functional lineage assignments (right) for predicted HGT-associated genes. Flow widths correspond to gene counts. The majority of predicted HGT genes map to Gammaproteobacteria and Bacilli, with smaller contributions from Actinomycetes and Chitinophagia, indicating predominant short-range horizontal gene exchange within bacterial lineages.

**Figure S3.** **Proteome Metabolic Pathway Annotation and PET-relevant functional roles across consortium members expanded to narrower categories.** Functional roles of consortium members were examined using proteome functional annotation. KEGG Orthology (KO) assignments and general protein functions were mapped to KEGG pathways **(A)** while proteins not assigned core metabolic pathways and only enzyme classifications or unassigned KO, were sorted and functional predictions were assigned into broad categories based on gene annotation and function within each strain **(B)**. Categories include hydrolytic/oxidative enzymes, redox handling, ethylene glycol/dehydrogenation, outer membrane and aromatic transport, protein folding, sorting, degradation, biofilm and quorum sensing, growth and regulation, amino acid metabolism, and stress and starvation. Bars represent the proportional contribution of each functional category per strain, highlighting strain-specific functional biases and a division of labor within the consortium during PET colonization, depolymerization, and downstream substrate assimilation.

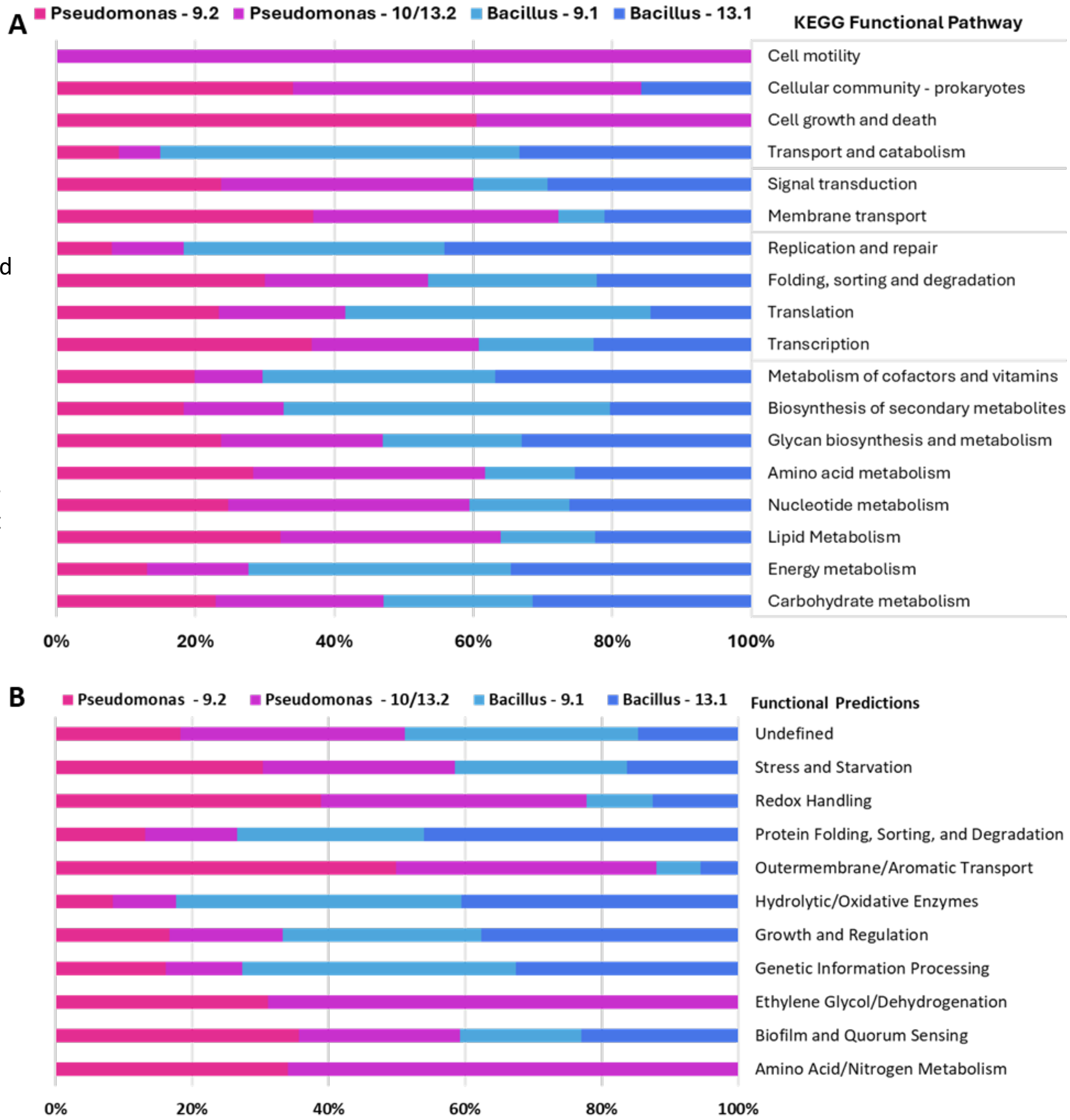

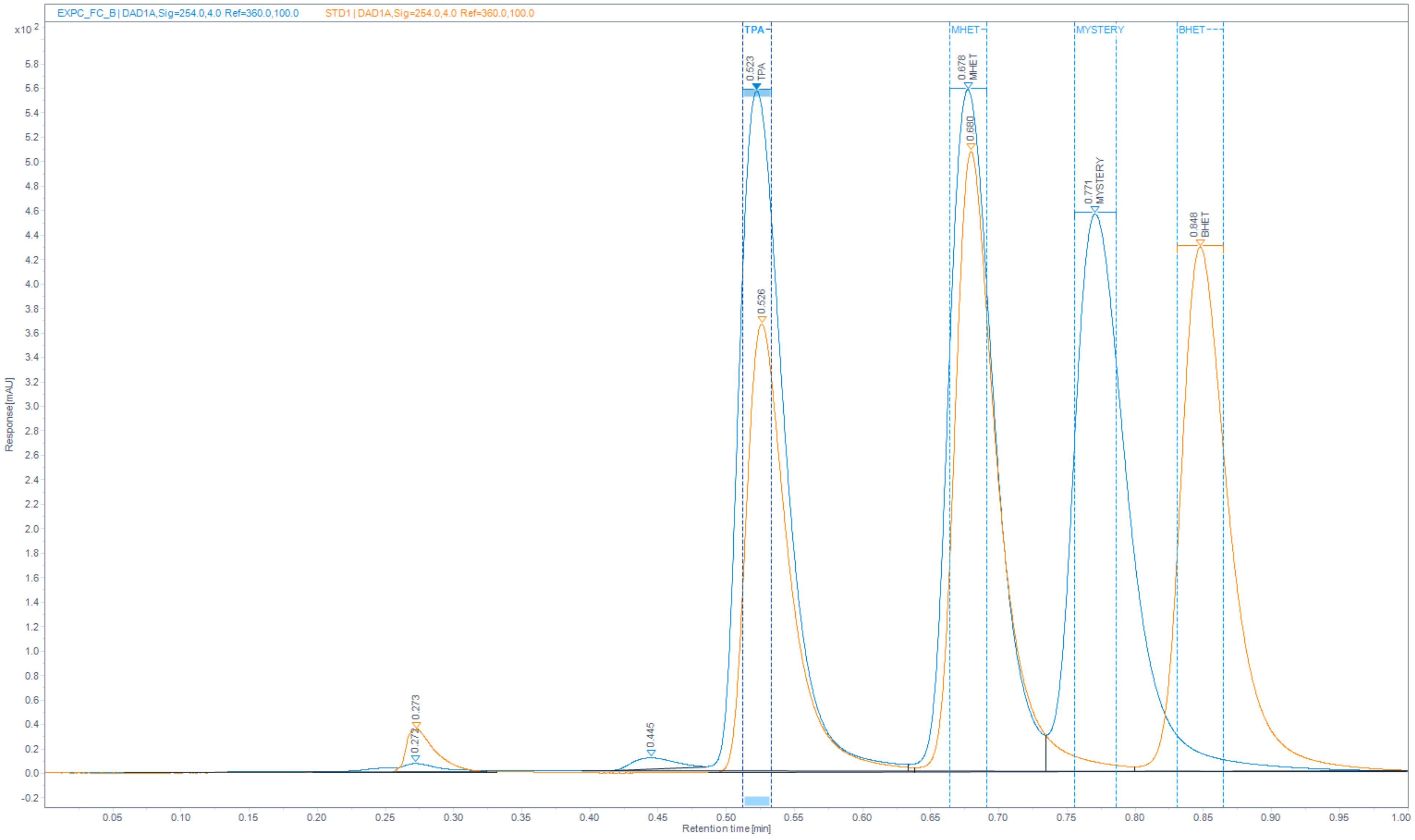


**Figure S4. HPLC chromatograms of consortium incubated with MHET overlaid with combined standards containing TPA, MHET and BHET show 'Mystery' Compound.** Retention times (in minutes) for aromatic monomers TPA and the oligomers MHET and BHET were found to be at 0.52, 0.68 and 0.85, respectively. A compound observed when incubated with MHET solely, shows a 'mystery' compound at 0.77 minutes, distinct from any canonical PETase byproducts.

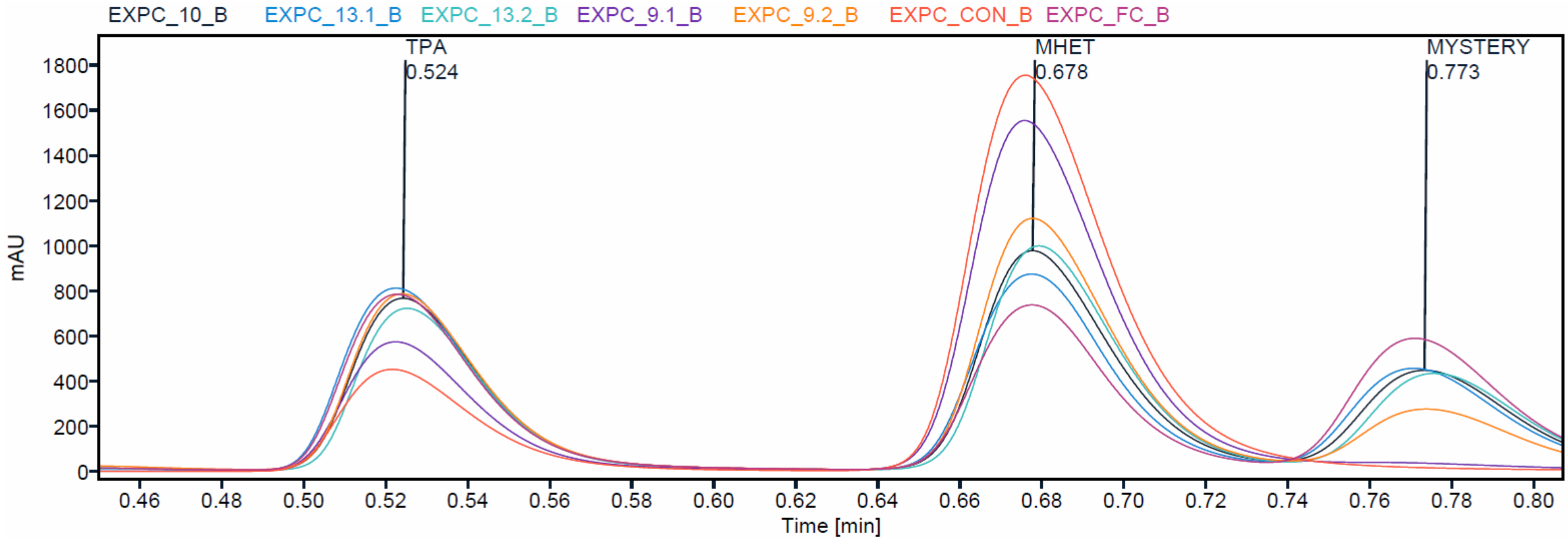


**Figure S5. HPLC chromatogram overlay of all strains compared with abiotic control (CON) and the full consortium (FC).** The abiotic control (CON) shown in dark orange shows a small amount of autohydrolysis of MHET to TPA. *Bacillus* strains, 9.1 in purple and 13.1 in blue. *Pseudomonas* strains are shown in light orange (9.2), black (10), and green (13.2). All five strains combined are shown in burgundy (FC). The presence of the mystery compound peaks in the FC and does not appear to be present when *Bacillus* strain 9.1 is incubated with MHET. Varying hydrolysis and conversion rates between strains indicates individual strain handling of MHET differs.

**Table S1: Distribution of DarkHorse consortium genes and their lineage probability index (LPI) score**

| *Bacillus* | | | | | | *Pseudomonas* | | | | | |
|---|---|---|---|---|---|---|---|---|---|---|---|
| **9.1 HGT Breakdown** | | | **13.1 HGT Breakdown** | | | **9.2 HGT Breakdown** | | | **10/13.2 HGT Breakdown** | | |
| | Count | LPI (min) | | Count | LPI (min) | | Count | LPI (min) | | Count | LPI (min) |
| **Viruses** | **14** | **0.003** | **Viruses** | **115** | **0.016** | **Viruses** | **11** | **0.002** | **Viruses** | **6** | **0.002** |
| Heunggongvirae | 14 | 0.003 | Heunggongvirae | 115 | 0.016 | Heunggongvirae | 11 | 0.002 | Heunggongvirae | 6 | 0.002 |
| **Eukaryota** | **2** | **0.105** | | | | **Eukaryota** | **32** | **0.004** | **Eukaryota** | **11** | **0.022** |
| Metazoa | 2 | 0.105 | | | | Fungi | 2 | 0.004 | Fungi | 1 | 0.022 |
| | | | | | | Metazoa | 3 | 0.006 | Metazoa | **8** | **0.118** |
| | | | | | | | | | Viridiplantae | **3** | **0.024** |
| **Bacteria** | **5008** | **0.407** | **Bacteria** | **4756** | **0.398** | **Bacteria** | **5388** | **0.418** | **Bacteria** | **4528** | **0.422** |
| | | | Thermodesulfobacteriota | 1 | 0.421 | Cyanobacteriota | 2 | 0.418 | Nitrospirota | 2 | 0.455 |
| | | | Spirochaetota | 1 | 0.421 | Campylobacterota | 2 | 0.418 | Cyanobacteriota | 2 | 0.424 |
| | | | Cyanobacteriota | 2 | 0.398 | Mycoplasmatota | 2 | 0.418 | Bacteroidota | 26 | 0.422 |
| | | | Myxococcota | 2 | 0.398 | Bacteroidota | 7 | 0.418 | Actinomycetota | 75 | 0.426 |
| Bacteroidota | 2 | 0.42 | Bacteroidota | 3 | 0.421 | Actinomycetota | 18 | 0.419 | Bacillota | 149 | 0.427 |
| Actinomycetota | 3 | 0.42 | Actinomycetota | 7 | 0.421 | Bacillota | 15 | 0.419 | Pseudomonadota | 4274 | 0.591 |
| Campylobacterota | 6 | 0.42 | Campylobacterota | 10 | 0.421 | Pseudomonadota | 5342 | 0.593 | | | |
| Pseudomonadota | 260 | 0.407 | Pseudomonadota | 258 | 0.408 | | | | | | |
| Bacillota | 4737 | 0.617 | Bacillota | 4472 | 0.618 | | | | | | |

For each consortium member, all predicted genes were assigned a lineage probability index (LPI) score using DarkHorse to assess potential horizontal gene transfer. Distributions show the frequency of genes across LPI values for each genome. Empirically determined thresholds of LPI ≤ 0.6 (broad cutoff) and LPI ≤ 0.4 (stringent cutoff) were used to identify candidate horizontally transferred genes. Lower LPI scores indicate reduced lineage consistency and a higher likelihood of horizontal acquisition. 166 exhibited very low LPI scores (<0.2), suggesting the strongest signal of horizontal inheritance. False positives were identified as Eukaryotic matches with low sequence coverage and phage derived genes. 1128 candidates were detected within all five organisms.

**Table S2:** **DarkHorse HGT Candidates Sorted by Function**

| Primary Category | Sub-Category | Protein Name / Family | Function |
|---|---|---|---|
| **Secretion, Transport & Membrane** | *Secretion Systems* | Type VI Secretion Protein | Involved in bacterial secretion. |
| | | Twin-Arginine Translocase (TatA/TatE) | Facilitates secretion of folded proteins. |
| | *Membrane & Envelope* | Cell Wall-Binding Protein | May aid in cell envelope stability. |
| | | Exosporium Protein ExsB | Structural component of bacterial spores. |
| | | Curlin | Associated with biofilm formation. |
| | | Adhesins | Aid in bacterial attachment to surfaces. |
| | *ABC Transporters* | Nickel, Peptide, Glycine, Methionine, etc. | Facilitate uptake of essential nutrients. |
| | | Putrescine Permease (PotI) | Involved in polyamine uptake. |
| | | Benzoate Transporter | Assists in movement of organic compounds. |
| | *Other Transporters* | MFS Transporters | Mediate sugar and metabolite transport. |
| | | Sodium:Solute Symporter | Uses sodium gradient for transport. |
| | | Iron Transporters (FeoB, FetB, etc.) | Involved in iron homeostasis. |
| | | OprD Family Porin | Functions in selective small molecule transport. |
| | *Protein Handling* | Clp Protease | Functions in protein degradation and stress response. |
| | | Protein-Disulfide Isomerase | Aids in protein folding. |
| **DNA Processing** | *Replication* | DNA Replication Protein | Essential for genome duplication. |
| | | Single-Stranded DNA-Binding Protein | Stabilizes single-stranded DNA. |
| | | DnaB-Like Helicase | Involved in DNA unwinding during replication. |
| | *Recombination* | Site-Specific Integrase | Integrates mobile elements into chromosomes. |
| | | Recombinase Family Protein | Facilitates DNA recombination and repair. |
| | | RuvA (Holliday Junction) | Assists in DNA recombination and repair. |
| | *Packaging & Modification* | DNA Packaging Protein | Assists in genome encapsulation. |
| | | Exodeoxyribonuclease VII Small Subunit | DNA processing enzyme. |
| | | HNH Endonuclease | Cuts DNA at specific sequences for recombination. |
| | | Metal-Dependent Hydrolase | Functions in nucleic acid metabolism. |
| | | Uracil-DNA Glycosylase | DNA repair enzyme. |
| **Gene Expression** | *Regulators* | Sigma Factors (70, 54, Sporulation) | Regulate bacterial transcription. |
| | | Sigma4 Domain Protein | A regulatory factor in gene expression. |
| | | Cro/Cl Family | Controls lytic/lysogenic switch. |
| | *Regulatory Families* | MarR, FadR, LysR, TetR, XRE | Control stress, metabolism, and antibiotic resistance. |
| | | AraC, GntR, LytR, TrmB, YeiL, Fis | Regulate metabolic and environmental response genes. |
| | | DNA-Binding Response Regulators | Control two-component bacterial signaling. |
| | *Transcription* | Antitermination Protein Q | Prevents premature transcription termination. |
| **Metabolism** | *Carbohydrate & Energy* | Beta-Galactosidase | Breaks down lactose and other sugars. |
| | | D-Ribose Pyranase | Sugar metabolism enzyme. |
| | | Thiamine Phosphate Synthase | Required for thiamine biosynthesis. |
| | | Phosphate Acyltransferase | Functions in fatty acid metabolism. |
| | | Phosphate Butyryltransferase | Involved in fermentation pathways. |
| | *Amino Acid* | Peptide Synthetase | Facilitates peptide biosynthesis. |
| | | Peptidases (M16, M24, etc.) | Enzymes involved in protein degradation. |
| | | Glutamine ABC Permease | Aids in nitrogen metabolism. |

| | | | |
|---|---|---|---|
| | *Nucleotide & RNA* | UMP Kinase | Participates in nucleotide metabolism. |
| | | Ribonuclease III | RNA processing enzyme. |
| | | RNA Polymerase Subunit Sigma-70 | Regulates transcription initiation. |
| | | 50S Ribosomal L3 Methyltransferase | Involved in ribosomal function. |
| | | Elongation Factor P (EF-P) | Facilitates bacterial protein synthesis. |
| | *Lipid & Coenzyme* | Acetyl-CoA BCCP Subunit | Essential for fatty acid biosynthesis. |
| | | Cobalamin Biosynthesis Protein CbiX | Involved in vitamin B12 production. |
| | | Molybdenum Cofactor Biosynthesis | Important for enzymatic redox reactions. |
| | *Oxidative / Redox* | Aldehyde Dehydrogenase (NADP+) | Metabolizes aldehydes in respiration. |
| | | GMC Family Oxidoreductase | Involved in oxidation-reduction reactions. |
| | | Xanthine Dehydrogenase | Functions in nitrogen metabolism. |
| | | 3-Oxoadipate Enol-Lactonase | Breaks down aromatic compounds. |
| | | Enoyl-CoA Hydratase | Participates in fatty acid degradation. |
| | | Diguanylate Cyclase | Controls signaling via cyclic di-GMP. |
| | *Respiratory* | Cytochrome B | Involved in electron transport chains. |
| | | NADH:Quinone Oxidoreductase | Essential for bacterial respiration. |
| | | Methyl-Accepting Chemotaxis Protein | Helps bacteria sense environmental signals. |
| **Stress & Defense** | *Environmental* | Universal Stress Protein | Helps bacteria survive under stressful conditions. |
| | | Cold-Shock Protein | Assists in adaptation to cold environments. |
| | *Resistance & Toxin* | Zeta Toxin Family | Involved in toxin-antitoxin systems. |
| | | BrnA Antitoxin Family | Counteracts bacterial toxins. |
| | | Superinfection Immunity Protein | Prevents infection by similar phages. |
| | | Bacterioferritin | Iron storage; protects against oxidative stress. |
| **Hypothetical** | *Unknown Domains* | DUF (1272, 2384, 2938, etc.) | Unknown functions; potentially metabolic/stress roles. |

**Table S3: Genomic regions of genetic island cassettes found in strain 10**

| Cluster | Start | End | Size_bp | Genes | Key gene_examples |
|---|---|---|---|---|---|
| **SOS / DNA damage** | 2502 | 8232 | 5731 | 4 | repressor LexA, protein ImuA, protein ImuB, error-prone DNA polymerase |
| **Benzoate cassette** | 59412 | 66964 | 7553 | 7 | benzoate/toluate 1,2-dioxygenase subunit alpha, benzoate/toluate 1,2-dioxygenase subunit beta, benzoate/toluate 1,2-dioxygenase reductase component, MFS transporter, AAHS family, benzoate transport protein, catechol 1,2-dioxygenase, benzoate membrane transport protein |
| **Phenylacetic acid (PAA) cassette** | 160653 | 164328 | 3676 | 5 | ring-1,2-phenylacetyl-CoA epoxidase subunit PaaE, ring-1,2-phenylacetyl-CoA epoxidase subunit PaaD, ring-1,2-phenylacetyl-CoA epoxidase subunit PaaC, ring-1,2-phenylacetyl-CoA epoxidase subunit PaaB, ring-1,2-phenylacetyl-CoA epoxidase subunit PaaA |

**Table S4: LC–MS (ESI–) Summary of PET-Related Intermediates**

| **Compound** | **Molecular Formula** | **Neutral Exact Mass (Da)** | **[M–H]$^-$ *(m/z)*** | **Fragment Ions seen *(m/z)*** |
|---|---|---|---|---|
| **TPA** | $C_8H_6O_4$ | 166.0266 | **165.019** | 165, 91, 211* |
| **MHET** | $C_{10}H_{10}O_5$ | 210.0528 | **209.045** | 209, 91, 255* |
| **MMHET** (O-methyl-MHET) | $C_{11}H_{12}O_5$ | 224.0685 | **223.061** | 223+, 91, 269* |

*Fomate Adduct seen from mobile phase formic acid

+14 *m/z* due to added methyl group
